# Magnetic resonance imaging does not reveal structural alterations in the brain of synesthetes

**DOI:** 10.1101/196865

**Authors:** M. Dojat, F. Pizzagalli, JM Hupé

**Affiliations:** Univ. Grenoble Alpes, INSERM, CHU Grenoble Alpes, GIN, F-38000, Grenoble, France; Univ. Toulouse Paul Sabatier, CNRS, CerCo, F-31300, Toulouse, France

**Author notes:** Corresponding author: Michel Dojat, Grenoble Institut des Neurosciences, INSERM U1216, Bâtiment Edmond Safra, Chemin Fortuné Ferrini, 38700 La Tronche.

**Keywords:** MRI, Voxel-Based Morphometry, Volumetry, Sulci, Connectivity

## Abstract

Several publications have reported structural changes in the brain of synesthetes compared to controls, either local differences or differences in connectivity. In the present study, we pursued this quest for structural brain differences that might support the subjective experience of synaesthesia. In particular, for the first time in this field, we investigated brain folding in comparing 45 sulcal shapes in each hemisphere of control and grapheme-color synesthete populations. To overcome flaws relative to data interpretation based only on p-values, common in the synesthesia literature, we report confidence intervals of effect sizes. Moreover, our statistical maps are displayed without introducing the classical, but misleading, p-value level threshold. We adopt such a methodological procedure to facilitate appropriate data interpretation and promote the New Statistics approach. Based on structural or diffusion magnetic resonance imaging data, we did not find any strong cerebral anomaly, in sulci, tissue volume, tissue density or fiber organization that could support synesthetic color experience. Finally, by sharing our complete datasets, we strongly support the multi-center construction of a sufficient large dataset repository for detecting, if any, subtle brain differences that may help understanding how a subjective experience, such as synesthesia, is mentally constructed.

## 1. Introduction

Various synesthetic experiences are reported by a potentially underestimated fraction, possibly up to about 20%, of the healthy population (1). Synesthetes experience additional, systematic, arbitrary and involuntary associations. For instance, they may associate a specific color to some “graphemes” (the visual form of numbers or letters). To understand this subjective experience, we may hypothesize that some structural differences exist in the brain of synesthetes compared to controls that trigger and support these associations. In particular, the cross-activation theory considers increased connectivity between proximal regions; for grapheme-color association between areas involved in color perception and areas involved in grapheme recognition (2). Several studies have searched for such extra-numerous connections using structural Magnetic Resonance (MR) images or Diffusion Tensor Imaging (DTI) (see (3, 4) for reviews). Indeed, structural imaging coupled with Voxel Based Morphometry (VBM) method (5, 6) appears as a powerful tool for detecting possible differences in the brain of synesthetes compared to controls. VBM assesses local (voxel-by-voxel) differences between populations of interest in tissues volume, mainly grey matter (GM) and white matter (WM), (7-11). With DTI one can measure fractional anisotropy (FA) and then track again voxel-by-voxel possible differences of structural connectivity in synesthetes’ brains (9, 12-15). However, despites many reports of those, no structural differences might exist in the brain of synesthetes. Indeed, the careful examination of the reported differences concerning 19 studies on morphometry (n=11) and structural connectivity (n=8), indicates that no clear view emerges from the literature and suggests that the observed differences are false positives due to methodological issues (Hupé and Dojat, 2015).

The present paper pursues this quest for structural differences in the brain of synesthetes. Firstly, we replicated our previously published study (8) on a new, larger, population of grapheme-color synesthetes. We obtained different results from those previously reported 1) in re-analyzing our initial data with an updated version of the initial VBM pipeline used, 2) in analyzing the new synesthete and control populations data and 3) in pooling the two sets of data. Second, to complement this morphometric analysis, we 4) searched for possible structural connectivity differences based on Mean Diffusivity (MD) and FA extracted from DTI data. Finally, and for the first time on a population of synesthetes, we 5) explored the possible anatomical differences at the level of the sulci architecture using a sulcal-based morphometry approach. In order to prevent from the severe flaws of null-hypothesis significance testing (NSHT) (16) and the difficult control of false positives (17), we adopted in this paper, the new statistic approach (18) reporting effect sizes (ES) and confidence intervals (CI). Moreover, statistical maps were displayed without introducing the classical p-value level threshold, limiting data interpretation bias (19). None of our analyses allowed us differencing synesthete brains from control brains, based on either standard tissue volume, tissue density or fiber organization criteria, or newly introduced brain sulci descriptors. Following these analyses investigating different brain features, we conclude that a larger sample size for appropriate statistical power is mandatory to reveal if any structural differences exist in the brain of synesthetes compared to controls. Multi-center data sharing is the realistic means to increase the synesthete population size to study. To launch such a data sharing process, we render our data publicly available (see Discussion Section).

## 2. Materials and Methods

### 2.1 Data

We conducted two studies involving synesthetes, one during the period from 2010 to 2011, Study 1, described in (8) and a more recent study, Study 2. These studies were performed following project approval by the Institutional Review Board of Grenoble and written consent from the subjects. All subjects had normal color perception on the Lanthony D-15 desaturated color test (Richmond products). Synesthetic associations were strictly controlled and consistency checked using a modified version of the Synaesthesia Battery test (20).

#### 2.1.1 Study 1

Ten grapheme-color synesthestes and twenty-five controls participated in this study. For each subject, we acquired structural images on a Bruker 3T Medspec S300 whole body scanner equipped with a birdcage head coil using a T1-weighted 3D MP-RAGE image consisting in 176 sagittal partitions in two segments with an image matrix of 256х112 (read х phase). Further imaging sequence parameters were: TR/TE/TI: 16/4.96/903 ms, excitation pulse angle: 8°, acquisition matrix: 176х224х256 (x,y,z), fast phase encoding in anterio-posterior direction (112 steps per RAGE train, 2 segments), slow phase encoding in left-right direction, isotropic nominal resolution: 1mm, BW=130Hz/Px, readout in caudo-cranial direction, number of averages: 1 and total measurement time: 14min40s.

#### 2.1.2 Study 2

Twenty-one grapheme-color synesthetes and twenty-six controls participated in this study. For each subject we acquired a high-resolution structural MP-RAGE image on a 3T Philips Intera Achieva, using a 32 channels coil, 180 sagittal slices of 256х240 (read x phase). Further imaging sequence parameters were: TR/TE/TI: 25/3.7/800 ms, excitation pulse angle: 15°, isotropic nominal resolution: 1mm, BW=191Hz/Px, readout in anterio-posterior direction, number of averages: 1, sense factor anterio-posterior: 2.2, right-left: 2 and total measurement time: 9min41s. We also acquired diffusion-weighted images (DTI) using a DWI sequence (120x120 matrix size with 70 contiguous transverse slices, FOV= 240mm, 2x2x1.75 mm^3^ spatial resolution, TR= 6845 ms, TE=667 ms, FA= 90°, Sense Factor=2 to improve the signal to noise ratio, total acquisition time: 18 min). The encoding protocol included 60 different non-collinear directions (gradient factor b=1000 s/mm^2^), and one image without diffusion weighting used as reference volume.

For both studies, image acquisitions were performed at Grenoble MR imaging facility IRMaGe.

### 2.2 Data processing of structural images

#### 2.2.1 Voxel Based Morphometry (VBM)

We analyzed the structural images using the VBM approach (5) where region-wise volumetric comparison among groups of subjects is performed. It requires all individual images to be registered in the same space and segmented in different tissue classes. Then, registration parameters and a mixture of Gaussian distributions for brain tissue modeling have to be estimated. A generative approach has been proposed for such a model parameter estimation that alternates among classification, bias correction and registration (21). We used the data processing pipeline VBM8 (http://www.neuro.uni-jena.de/vbm/) that implements this approach as a toolbox extension of SPM8 (http://www.fil.ion.ucl.ac.uk/spm/software/spm8/) running with Matlab language. Compared to the previous one, this new version mainly improves the segmentation step in removing noise (22), estimating partial volume effects (23), taking into account local intensity variations and ensuring local coherency of tissue labels (24). A joint segmentation-registration approach is used to segment and warp the individual tissue probability maps into a common study-specific reference space. Then an affine registration may be applied for transformation into the Montreal Neurological Institute (MNI) referential space. Each structural image is segmented by attributing to each voxel a probability of being in white matter (WM), grey matter (GM), and cerebrospinal fluid (CSF). This procedure uses a maximum *a posteriori* estimation to take into account local variations of intensity and estimates a mixture model composed of several Gaussian distributions notably for pure tissue (3 gaussians) and partial volume effects (2 gaussians). To impose local coherence, a markovian approach introduces spatial prior into the model estimation. The high-dimensional DARTEL (21) registration algorithm iteratively computes deformation fields for warping each individual image to the common space. To counterbalance local deformations, expansion, or contraction, induced by highly non-linear registration and affine transformation, the tissues’ probability values computed were scaled by the Jacobian determinants of the deformations (“modulation step” (25)). Finally, following the recommendations of (26) for decreasing the false positive rate, we smoothed these “modulated” tissues probability maps using a 12-mm full-width at half-maximum Gaussian kernel (same pattern of results with an 8-mm kernel). A visual control of the sample homogeneity as implemented in VBM8 was realized based on the covariance between the images. No outlier was detected.

#### 2.2.2. Diffusion Tensor Imaging

DTI images were first denoised (27) and preprocessed using FSL software (http://www.fmrib.ox.ac.uk/fsl/). The images were corrected for geometric distortions caused by Eddy currents and intensity inhomogeneity. The diffusion tensor was estimated, and the local diffusion fractional anisotropy (FA) and mean diffusivity (MD) parameters were calculated for the entire brain in each participant. These parameters were computed from the three estimated eigenvalues describing the water diffusion in three orthogonal directions. FA represents the coefficient of variation of these eigenvalues interpreted as the directionality of water diffusivity into fibers (coherence). MD is the mean of the eigenvalues. These parameters are myelination markers, serving as measures of tissue density for MD and fiber organization for FA. For each participant, the non-diffusion weighted image (T2-weighted) was realigned to the corresponding structural image. The computed realignment parameters were applied to the corresponding MD volumes to be aligned to the structural image. Then they were warped using the deformation field previously computed for this image for warping to the common space, scaled by the Jacobian determinants of the deformations and smoothed (12 mm FWHM). Subsequently, MD volumes were analyzed on a voxel-by-voxel basis similarly to VBM-GM/WM volumes analysis (28). Because such a voxel-based analysis does not ensure a proper alignment of individual fiber tracts, a Track-Based Spatial Statistics (TBSS) was also processed (29). Individual FA data were nonlinearly realigned and a mean FA image was computed and used to define a mean FA skeleton on which all individuals FA data were projected.

#### 2.2.3 Sulci extraction and morphometry

“Sulcus-based morphometry” provides measures of the cortical fissures of the brain, which have been found to be associated with brain maturation (30), brain alteration with age or pathology (31, 32) and correlated in a population of twins (33). We investigated sulci morphometry in controls and synesthetes based on shape descriptors (width, length, mean depth, and total surface area) analyzed for each sulcus. We used Freesurfer (https://surfer.nmr.mgh.harvard.edu) to classify grey and white matter tissues and Morphologist 2013, an image-processing pipeline included in BrainVISA (http://brainvisa.info/web/index.html), to quantify the sulcal descriptors. Briefly, the Morphologist 2013 segmentation pipeline computes a brain mask, imports brain tissues (gray, white matter and CSF) classified by Freesurfer, performs gray/white surface identification and spherical triangulation of the external cortical surface of both hemispheres. Sulci were then automatically labeled according to a predefined anatomical nomenclature (34). The number of voxels on the junction between the sulcal mesh and the brain hull gives a measure of sulcal length. The mean depth is defined as the mean of geodesic distances computed for all voxel belonging to the sulcal fundus, from the bottom of the sulcus to the brain hull, along the cortical mesh. The surface area is the total area of the sulcal mesh. The sulcal width is obtained by dividing the enclosed CSF volume by the sulcal surface area; see (35) for details. We considered that a sulcus could not be measured for a given subject if the four measures (length, width, surface and depth) were equal to zero (that was the case for most zero values). Such cases appeared when individual variability was too high compared to the reference population. The pattern recognition algorithm then failed to sulcus identification meaning that the corresponding sulcus was absent for this subject or not measurable. When a sulcus was not measured for more than 11% of subjects either in the control group or in the synesthete group (that was, in either 5 subjects in the control group or 3 subjects in the synesthete group), it was removed from the analysis (it does not make sense to compare different sulci if tested on too many different subjects). When a sulcus was missing for a subject, we attributed the median value of all other subjects (controls and synesthetes, left and right sides, independently) for that sulcus. The interpolated value was used only for the sum and the subsequent normalization. We normalized each measure by dividing the within subject sum of the sulci independently for length, depth, surface and width. We computed Pearson’s correlation between left and right sulci values to consider pooling the right and left values or not.

### 2.3 Statistical analysis

#### 2.3.1 Structural images

We compared the regional tissue probability maps (modulated and smoothed as described above) of controls and synesthetes by performing a voxel-wise univariate analysis using the general linear model (GLM) as implemented in SPM8. Because the global brain size can vary across subjects, we included brain volume as a factor of noninterest in our statistical tests. In order to calculate the global brain volume, we used the modulated images by summing together the GM and WM probabilities of all voxels. To avoid possible edge effects between different tissue types, we applied an absolute intensity threshold mask of 0.1 on each tissue probability. For Study 1, average age slightly differed between the 2 groups (29.8 vs. 36.4, years p= 0.082), and our synesthete group had more women (7/10 vs. 10/25 in our control group). Both factors may generate local differences not related to synesthesia, so we also included sex and age as factors of noninterest. In Study 2, the two groups were not different in term of age (28 vs 28.4 years) and sex (18/21 vs 19/26 in our control group). However, in order to compare the two studies, we adopted the same model including sex and age as cofactors at the expense of the number of degrees of freedom.

#### 2.3.2 Diffusion Tensor Imaging

For MD data, we performed (Study 2) a voxel-wise univariate analysis using a GLM model similar to that used for tissue analysis. For FA data, we compared FA projections onto the mean population skeleton between synesthetes and controls using voxel-wise permutation tests (36). We performed cluster-based statistics with the "threshold-free cluster enhancement" (TFCE) approach as implemented in FSL “randomize” function (37), without any spatial smoothing.

#### 2.3.3. Sulci

We compared the distribution of sulcal length, mean depth, surface and width between the two populations across both studies for right and left hemispheres. We computed 99.9% confidence intervals for the group difference using a linear model, with age and sex as covariates. We chose quite arbitrarily the 99.9% value to partially account for multiple comparisons (4 measures in 61 sulci) while avoiding being too conservative, because of possible correlations between measures (a 99.9% CI corresponds to a 95% family-wise CI for 50 independent comparisons). We could not perform multivariate analyses because of the large number of missing values (not all sulci could be identified in every subject).

#### 2.3.4 Data exploration

To compare with our results published in 2012 we considered the family wise error (FWE correction for multiple comparison) measured at the cluster level. We considered two regions of interest (spheres, radius 6 mm) located at the coordinates of the clusters found in (8) that survived a strict FWE correction, in the right retrosplenial cortex (RSC) and the left superior temporal sulcus (STS). Two additional regions in the fusiform gyrus and in the parietal cortex were considered. For each sphere, we computed in both studies WM local volume estimated using our general linear model for the two groups. Recently Eklund et al. (17) showed that such a multiple comparison correction procedure did not guarantee at all the control of false positive at a cluster level. Besides standard p-values for comparison with published studies, we reported effect sizes (ES) and confidence intervals (CI) to facilitate appropriate data interpretation in context. Our interpretation was anyway based on non-thresholded maps. For sulci analysis, no previous study was reported and then NHST was not considered. Moreover, to facilitate data interpretation, our statistical maps are displayed without introducing the classical p-value level threshold. We adopt the dual-coding approach proposed in (19) where differences in effect size (beta estimates) are color-coded and associated t-statistics mapped to color transparency.

## 3. Results

### 3.1 Reproducibility tests

#### 3.1.1 Study 1

We compared the local distributions of WM and GM in the brains of 10 synesthetes with the brains of 25 controls. We first used a cluster forming threshold at p<0.0001, similarly to (8) but we did not obtain any clusters of at least 70 voxels size. At a more lenient threshold (p<0.001) we found local increases of WM in synesthetes compared to controls in the right retrosplenial cortex (Figure 1A) and the left anterior middle temporal gyrus (Figure 1B), similarly to (8) (see their Figure 6). However, these clusters were found at a statistical threshold non-corrected for multiple comparisons (see Table 1). At this threshold new clusters appeared (see Table 1) in the right inferior parietal lobe (Figure 1C) and in the vicinity of color-sensitive regions of the fusiform gyrus (8, 38-40) (posterior cluster in Figure 1B). These two regions were close to some reported results (9-11). In coherence with the previously published analysis of these data (8) we did not find, at our statistical threshold (p<0.001 voxel-wise, cluster extent k > 70), any increase of WM for control subjects compared with synesthetes as well as no difference either way in GM.

**Figure 1.**
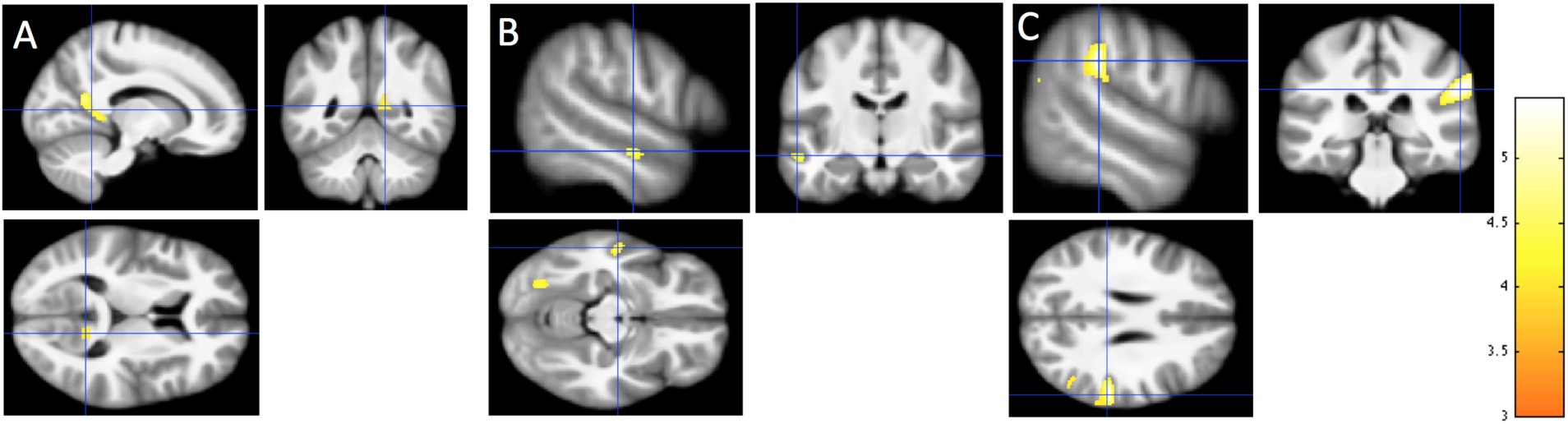
Local increases of WM in synesthetes compared with controls. Detected changes are projected onto the study-specific structural image transformed in the MNI space. (A) Increase in the right RSC (14, -57, 6). (B) Increase in the left anterior middle temporal gyrus (Blue cross: -56 -15 -14). Note the second cluster in the posterior part located in the fusiform gyrus (-26 -73 -12). (C) Increase in the right inferior parietal lobe (58 -31 28). All coordinates are in MNI space expressed in mm. Cluster-forming threshold p<0.001 uncorrected, t> 3.4, k > 70. T scale is between 3 and 5.5. Neurological convention (Right = Right).

**Table 1.** Local increase of WM in synesthetes (n=10) compared with controls (n=25)

|  | Cluster size (mm<sup>3</sup>) | x (mm) | y (mm) | z (mm) | Max T value | FWE-corr |
| --- | --- | --- | --- | --- | --- | --- |
| Right Inferior parietal lobe | 911 | 58 | -31 | 28 | 5.43 | 0.141 |
| Right RSC | 489 | 14 | -57 | 6 | 4.54 | 0.334 |
| Fusiform gyrus | 187 | -26 | -73 | -12 | 3.96 | 0.365 |
| Left STS | 86 | -56 | -15 | -14 | 4.10 | 0.779 |

Note: (x, y, z) = MNI coordinates of the center of each cluster. Max **T** value is the voxel maximum in the corresponding cluster. FWE-corr is the **P**-value corrected for the FWE at the

Figure 2 shows the set of structural differences without the introduction of a threshold.

**Figure 2.**
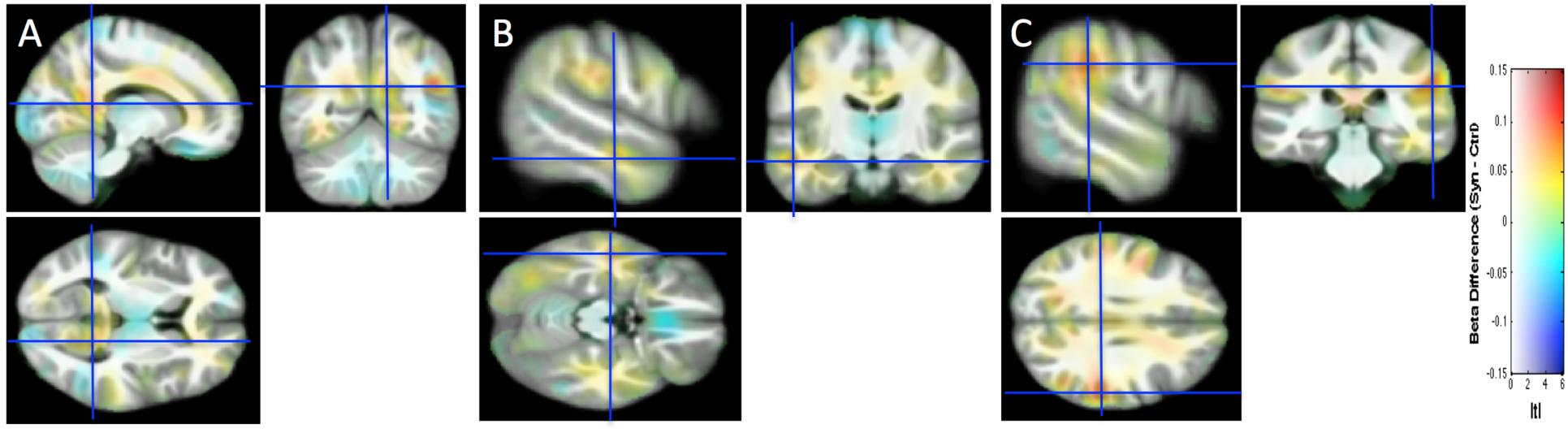
The structural differences between the two groups are projected without the introduction of a threshold onto the study-specific structural image transformed in the MNI space. Same slices, convention and cursor positions as in Figure 1. Beta difference scale is between -0.15 and 0.15 and mapped to color hue. T statistic magnitude is mapped to color transparency.

Figure 3 shows the individual data (denoted by circles for this first study) for the four regions of interest identified in Figure 2, using 6-mm spheres centered on the peak coordinates of Table I, as well as the 99.99% (corresponding to the arbitrary voxel-vise threshold of our statistical map used to identify clusters). Confidence Intervals (CI) of the group difference. By construction, the extent of each CI was expected to be above zero, since each value is the average of voxels with p-values < 0.001 for the group comparison. Note however that for small clusters (like in the left STS) this was not necessarily the case since the 6-mm sphere may extend beyond the cluster of small p-values voxels.

**Figure 3:**
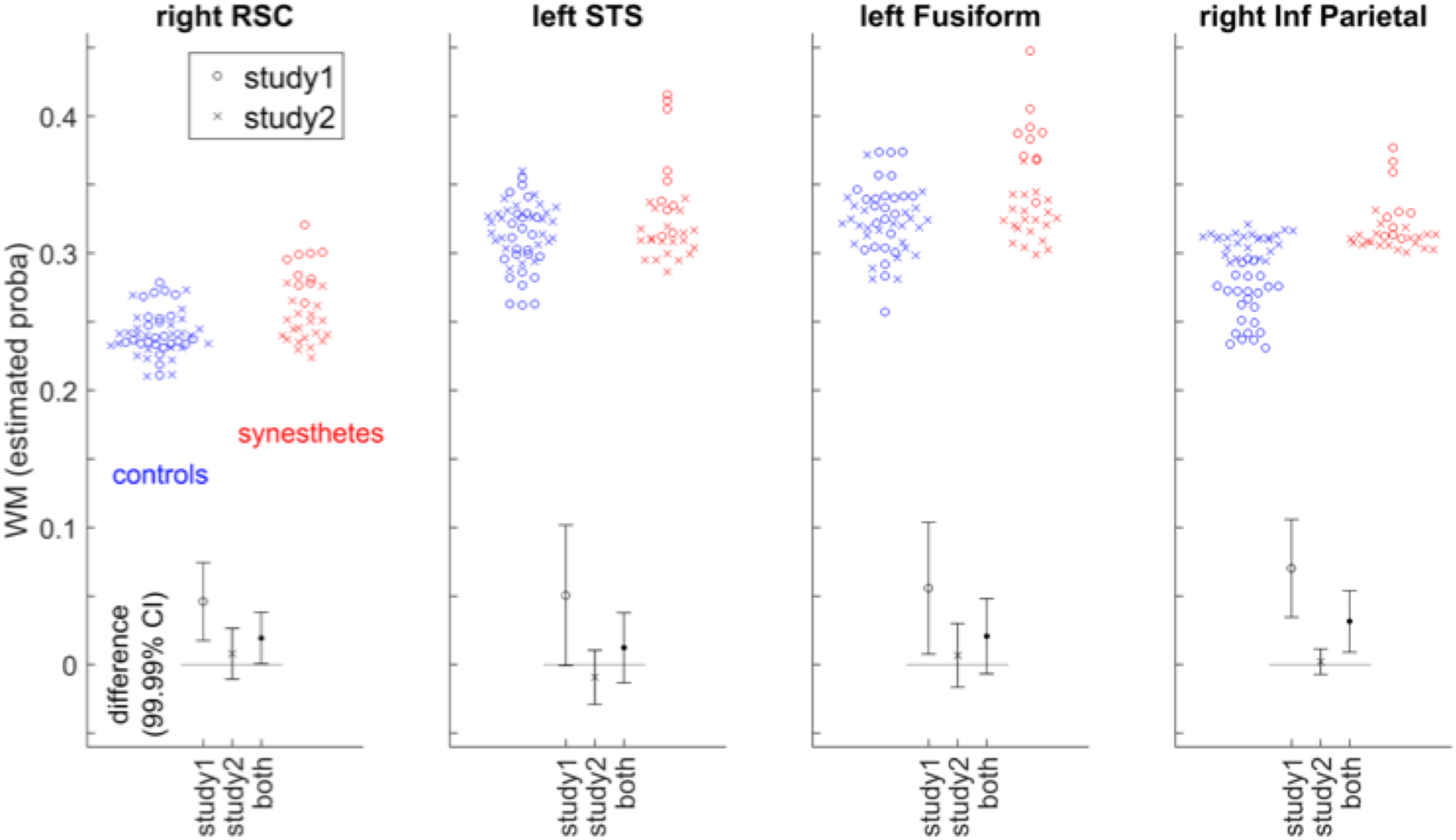
WM in synesthete and control groups, and 99.99% CI for the group difference, for four regions of interest centered in the right RSC, left STS, the right inferior parietal lobe and the left fusiform gyrus (coordinates in Table 1), for the subjects of study 1, study 2, and all subjects together. The WM tissue probability at each voxel was estimated by the linear model that included total brain volume, age and sex as covariates.

#### 3.1.2 Study 2

We compared the local distributions of WM and GM in the brains of 21 synesthetes with the brains of 26 controls. Figure 4 shows the structural differences of WM without introducing any arbitrary threshold, in the same slices as for Study 1. Figure 4 (crosses) indicates that structural differences are unlikely (or too small to be relevant and detectable) in the regions of interest defined in the first study (see also the CI of group differences for Study 2 in Figure 3). Using conventional 5% family-wise error risk, we did not find any ground (increase or decrease) for rejecting the Null hypothesis of no difference between the two groups. We obtained the same result for GM analysis. Following the protocol proposed by Kurth et al (41) we searched to assess whether the voxel-wise gray matter asymmetry in synesthetes group was significantly different from the voxel-wise gray matter asymmetry in control group No differences were found using a standard p<0.001 for individual voxels.

**Figure 4.**
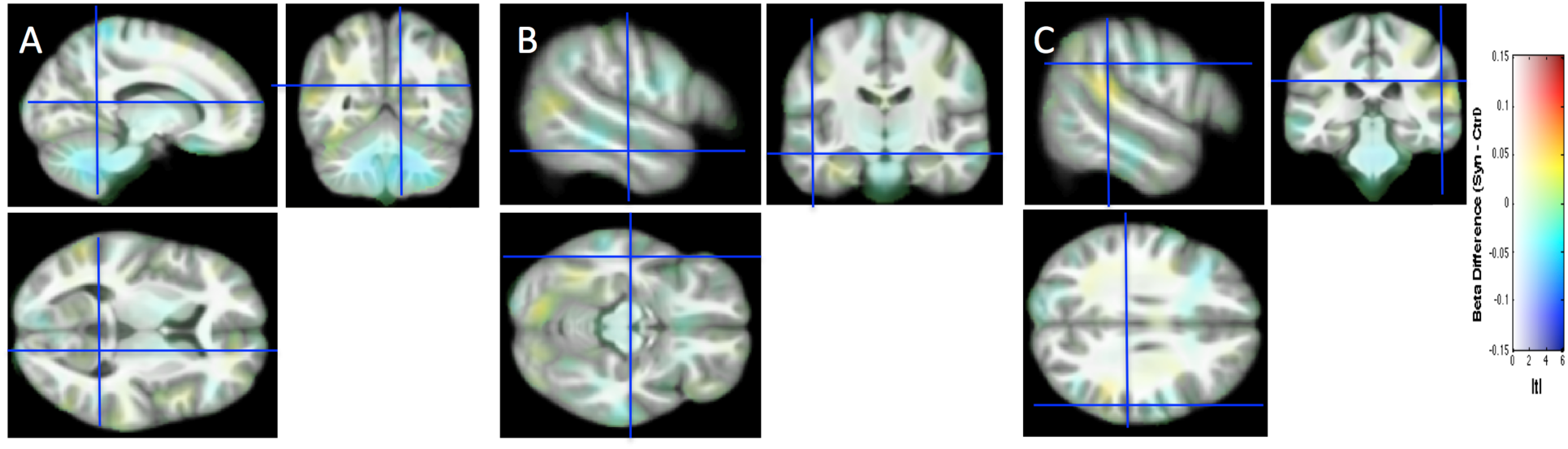
Study 2. Structural differences between the two groups are projected without the introduction of a threshold onto the study-specific structural image transformed in the MNI space. Same slices, convention and cursor position as in Figure 1.

#### 3.1.3 Pooling Data from Study 1 and Study 2

Here we considered pooling data from Study 1 and Study 2 (synesthetes = 31, controls = 51). Because data were acquired on two different scanners, 3T Bruker for Study 1 and 3T Philips for Study 2, we were particularly meticulous with the quality check after the preprocessing including realignment and segmentation steps. We used the module available in VBM8 for checking homogeneity of the segmented volumes GM and WM separately using covariance. We added in our general linear model a fourth covariable of non-interest corresponding to the two different conditions of data acquisition. Figure 5 shows all the WM differences without introducing any arbitrary threshold. Figure 3 shows the Confidence Intervals for the regions considered in Figures 2 and 4. Two clusters emerged at our threshold, none of them allowing us to reject the Null Hypothesis at the 0.05 FWE level (see Table2).

**Figure 5.**
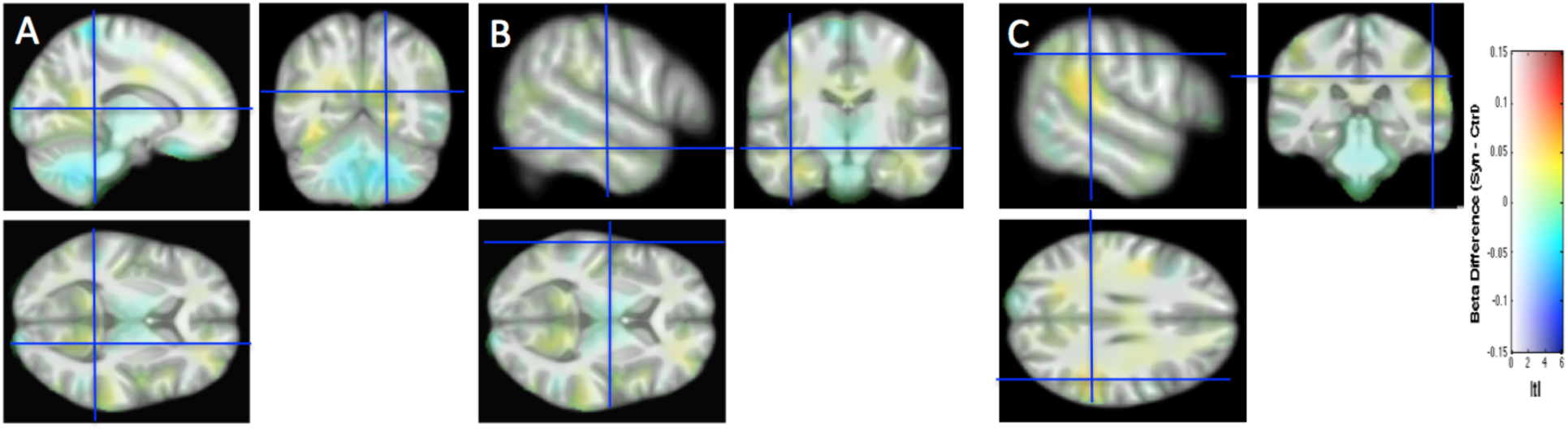
All the structural differences between the two groups are projected without the introduction of a threshold onto the study-specific structural image transformed in the MNI space. Same slices, convention and cursor position as in Figure 1.

In order to provide an idea of the power of these “Null” results, one may consider the upper (for increases) and lower (for decrease) limits of the CIs. For the increases in synesthetes relative to control reported in Figure 3, the largest differences of WM probability compatible with our measures in the 4 regions of interest were about 0.05 (based on a 99.99% CI). In the voxels with the maximal statistical differences (maxT = 4.39 and minT=-4.90) the upper and lower limits of the WM probability differences were respectively 0.0555 and -0.0525.

### 3.2 DTI

We compared the local distributions of MD in the brains of 21 synesthetes with the brains of 25 controls. We did not to find any “statistically significant’ difference using the using the p-value based procedure as done for WM and GM tissue (Cluster-forming threshold p<0.001 uncorrected, t> 3.4, k > 70). Figure 6 shows all the differences without the introduction of an arbitrary threshold at the same spatial coordinates we used for brain tissue comparison.

**Figure 6.**
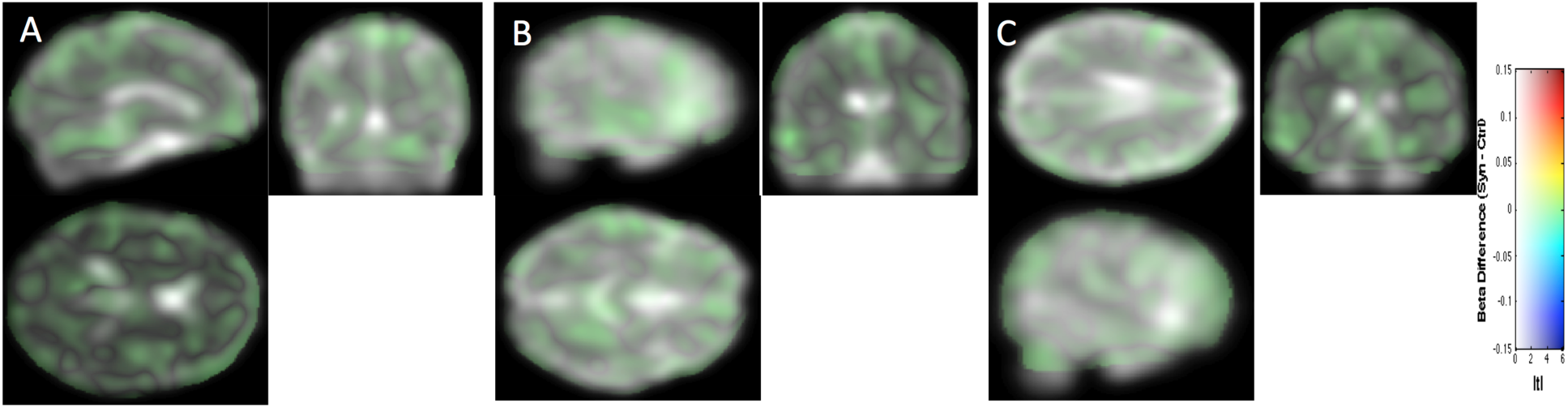
All the mean diffusivity differences between the two groups are projected without the introduction of a threshold onto the MD image of one subject transformed in the MNI space.

We also did not find any “statistically significant” difference of FA data, computed using permutation tests on TFCE cluster-based statistics (see Methods). An exploratory analysis (i.e. p<0.05 without correction for multiple comparisons) led to three clusters of at least 10 voxels of increase in FA, reported in Table 3, and decrease in FA, reported in Table 4, for synesthetes compared to controls. In average the detected tracks represented 9% (114599 voxels) of the brain volume.

**Table 2.** Local increase of WM in synesthetes (n=32) compared with nonsynesthetes (n=32)

|  | Cluster size (mm<sup>3</sup>) | x (mm) | y (mm) | z (mm) | Max T value | FWE-corr |
| --- | --- | --- | --- | --- | --- | --- |
| Right Superior temporal lobe | 1084 | 68 | -37 | 18 | 4.13 | 0.127 |
| Parahippocampal gyrus | 227 | -24 | -73 | -15 | 3.81 | 0.566 |

Note: (**x**, **y**, **z**) = MNI coordinates of the center of each cluster. Max **T** value is the voxel maximum in the corresponding cluster. FWE-corr is the **P**-value corrected for the FWE at the cluster level. We obtained these 2 clusters when thresholding **P**<0.0001 for individual voxels, with a minimum cluster size of 70 mm<sup>3</sup>.

**Table 3.** Local increase of FA in synesthetes (n=21) compared with controls (n=26)

|  | Cluster size<br>(mm<sup>3</sup>) | x<br>(mm) | y<br>(mm) | z<br>(mm) | Max T<br>value | FWE-corrected |
| --- | --- | --- | --- | --- | --- | --- |
| Right frontal cortex | 101 | 26 | 27 | 5 | 4.74 | uncorrected |
| Right caudate nucleus | 26 | 8 | -2 | 9 | 4.41 | uncorrected |
| Right post central gyrus | 10 | 26 | -34 | 56 | 4.11 | uncorrected |

Note: (x, y, z) = MNI coordinates of the center of each cluster. Max **T** value is the voxel maximum in the corresponding cluster. We obtained these 3 clusters with a non-parametric permutation testing analysis when thresholding $p < 0.05$ for individual voxels uncorrected for multiple comparison.

**Table 4.** Local decrease of FA in synesthetes (n=21) compared with controls (n=26)

|  | Cluster size<br>(mm<sup>3</sup>) | x<br>(mm) | y<br>(mm) | z<br>(mm) | Max T<br>value | FWE-corrected |
| --- | --- | --- | --- | --- | --- | --- |
| Middle frontal gyrus | 16 | 32 | 35 | 1 | 3.86 | uncorrected |
| Left superior frontal gyrus | 14 | -18 | 50 | 3 | 3.48 | uncorrected |
| Right inferior frontal gyrus | 14 | 34 | 35 | 5 | 3.8 | uncorrected |

Note: (x, y, z) = MNI coordinates of the center of each cluster. Max **T** value is the voxel maximum in the corresponding cluster. We obtained these 2 clusters with a non-parametric permutation testing analysis when thresholding $p < 0.05$ for individual voxels uncorrected for multiple comparison.

Figure 7 shows the difference when applying a threshold (p<0.05, panels A to C) and without the introduction of an arbitrary threshold (panel D). No clusters survived the application of a correction for multiple comparisons (TFCE) for p<0.05

**Figure 7.**
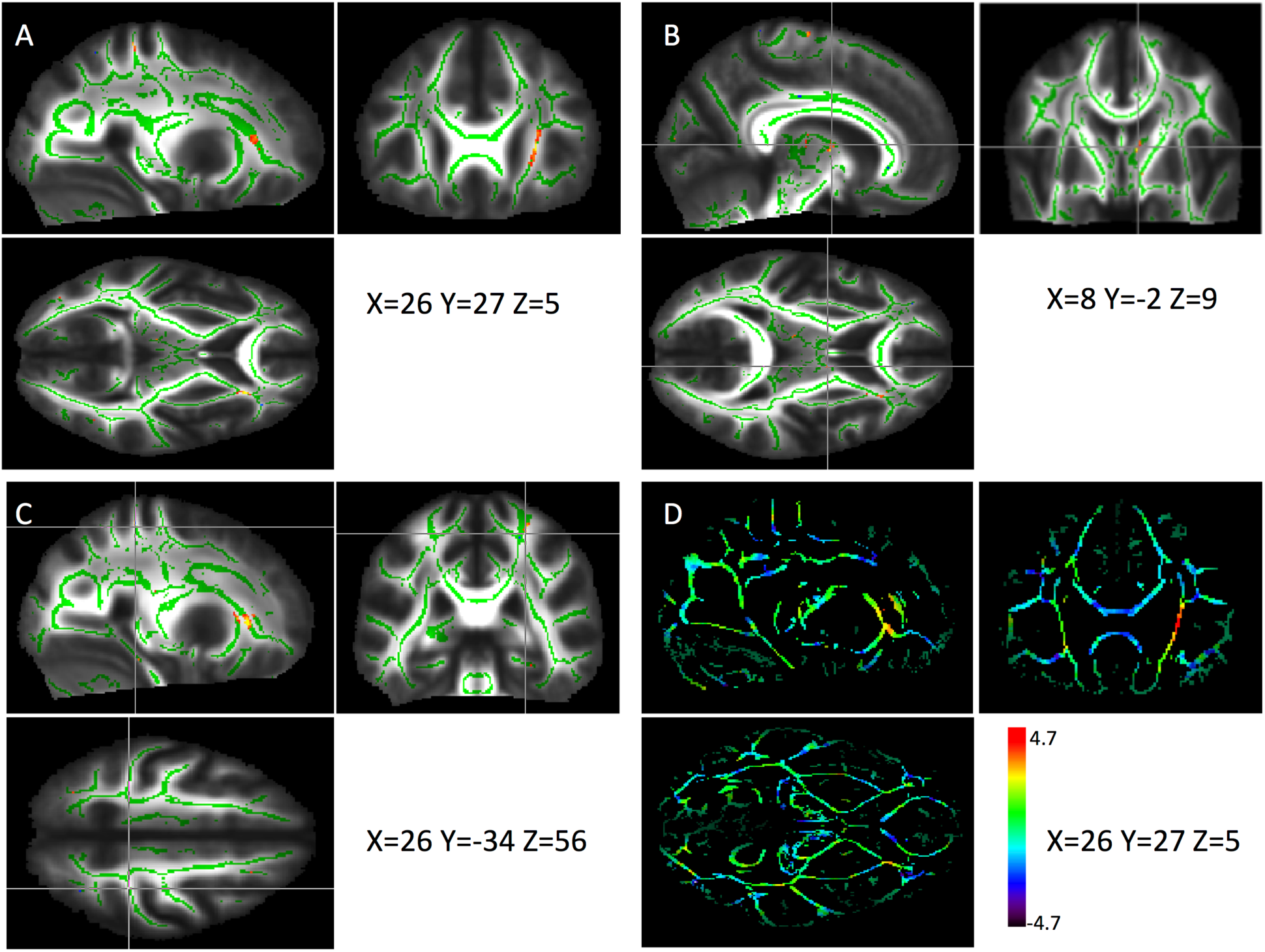
FA data. A, B, C: Increase in FA (in yellow-red) for synesthetes vs controls (p<0.05) projected onto the mean FA image (white matter skeleton in green). D: All FA differences between the two groups without the introduction of a threshold.

Figure 8 shows the distributions of FA mean values in the clusters identified in Tables 3 and 4, in controls and synesthetes, as well as the 95% Confidence Intervals of the group difference (no correction for multiple comparisons, age and sex were included as covariates).

**Figure 8:**
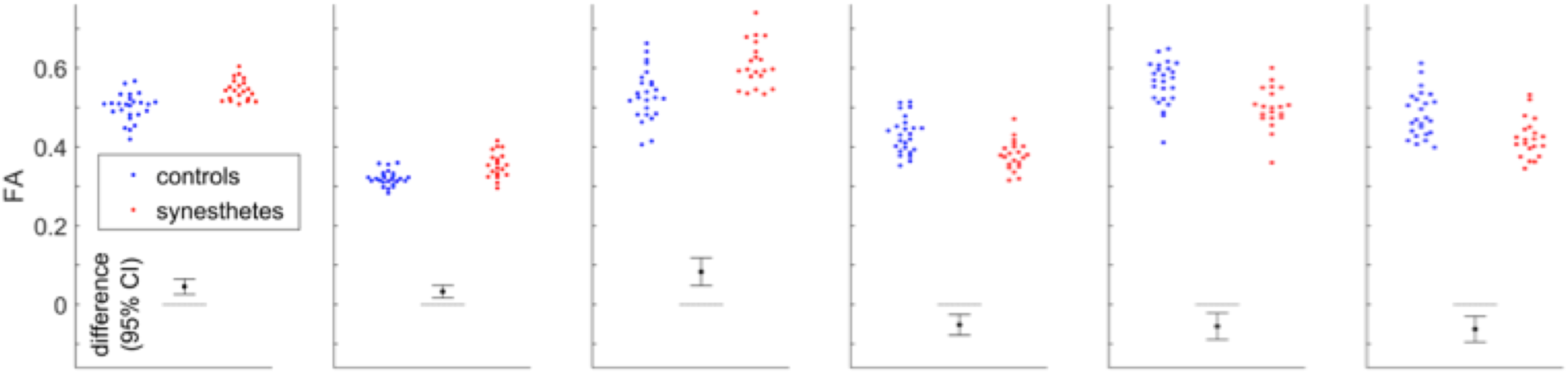
FA in synesthete and control groups, and 95% CI (no correction for multiple comparisons) for the group difference, for the six clusters identified in Tables 3 and 4 (same order); the first three graphs correspond to increases represented in red in Figures 7 A-C.

### 3.3 Sulci morphometry

Sixty-two sulci for the left hemisphere and sixty-one sulci for the right hemisphere were extracted for each individual for Study 1 and Study 2 (for the list of extracted sulci based on the Brainvisa sulci atlas see (34)). Our criterion on missing values (see methods) allowed us to have only up to 5/49 controls or up to 3/31 synesthetes with at least a missing sulcus. Sixteen sulci were then excluded (see Fig. 9). We normalized each value by dividing the within subject sum of the 90 sulci (45 right and 45 left), independently for length, depth, surface and width. Correlations (Pearson score) varied widely between corresponding left and right sulci, preventing from averaging the left and right values.

**Figure 9.**
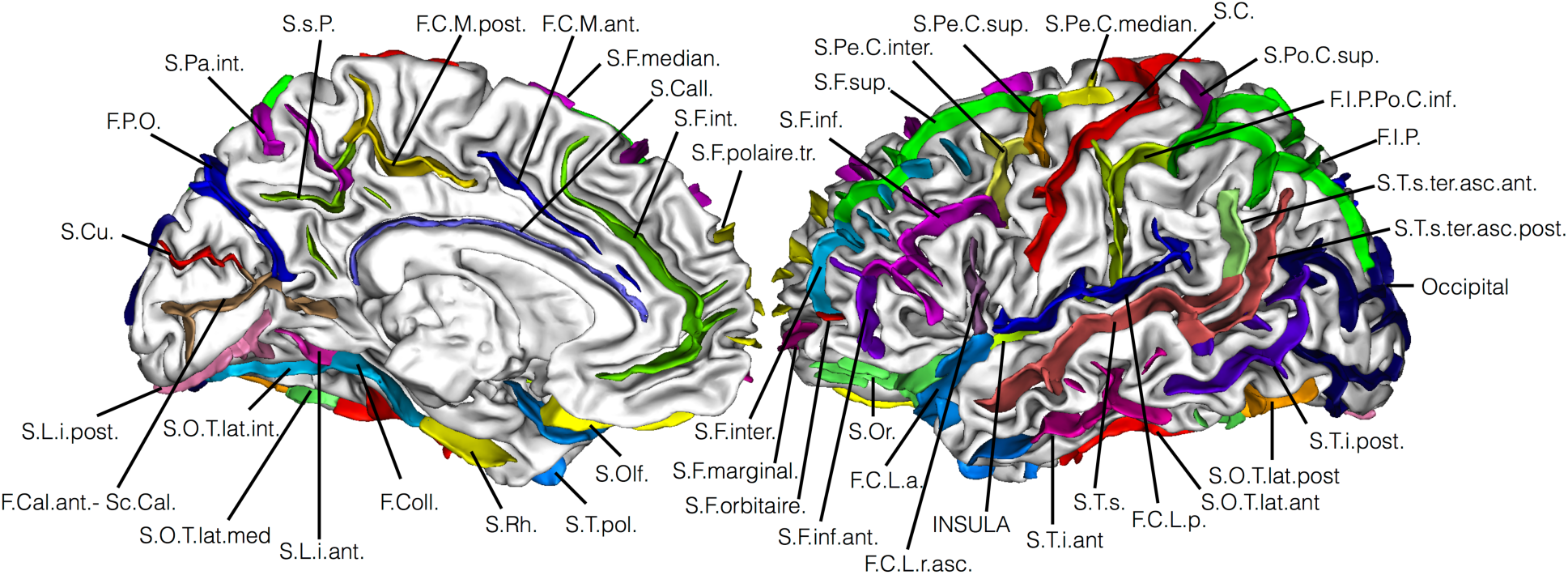
The 45 sulci we considered. F.C.L.a: anterior lateral fissure; F.C.L.p: posterior lateral fissure; F.C.L.r.asc: amending remus of the lateral fissure; F.C.M.ant.: calloso-marginal anterior fissure; F.C.M.post.: calloso-marginal posterior fissure; F.Cal.ant.-Sc.Cal.: calcarine fissure; F.Coll.: collateral fissure; F.I.P.: intraparietal sulcus; F.I.P.Po.C.inf.: sup. postcentral intraparietal sup. Sulcus; F.P.O.: parieto-occipital fissure; INSULA: Insula; Occipital: occipital lobe; S.C.: central sulcus; S.Call.: subcallosal sulcus; S.Cu.: cuneal sulcus; S.F.inf.: inferior frontal sulcus; S.F.inf.ant.: anterior inferior frontal sulcus; S.F.int.: internal frontal sulcus; S.F.inter.: intermediate frontal sulcus; S.F.marginal.: marginal frontal sulcus; S.F.median.: median frontal sulcus; S.F.orbitaire.: orbital frontal sulcus; S.F.polaire.tr.: polar frontal sulcus; S.F.sup.: superior frontal sulcus; S.Li.ant.: anterior intralingual sulcus; S.Li.post.: posterior intra-lingual sulcus; S.O.T.lat.ant.: anterior occipito-temporal lateral sulcus; S.O.T.lat.int.: internal occipito-temporal lateral sulcus; S.O.T.lat.med.: median occipito-temporal lateral sulcus; S.O.T.lat.post.: posterior occipito-temporal lateral sulcus; S.Olf.: olfactory sulcus; S.Or.:or bital sulcus; S.Pa.int.: internal parietal sulcus; S.Pe.C.inter.: intermediate precentral sulcus; S.Pe.C.median.: median precentral sulcus; S.Pe.C.sup.: superior precentral sulcus; S.Po.C.sup.: superior postcentral sulcus; S.Rh.:r hinal sulcus; S.s.P.: sub-parietal sulcus; S.T.i.ant.: anterior inferior temporal sulcus; S.T.i.post.: posterior inferior temporal sulcus; S.T.pol.: polar temporal sulcus; S.T.s.: superior temporal sulcus; S.T.s.ter.asc.ant.: anterior terminal ascending branch of the sup. temp. sulcus; S.T.s.ter.asc.post.: posterior terminal ascending branch of the sup. temp. sulcus.

Figure 10 shows t-scores for the differences between the two groups for our four measures. Similarly to Figure 5, we adopt a color scale that would highlight with salient colors (red and blue) only regions with potentially interesting differences, i.e. when the whole CI would be away from zero by at least one standard error above the mean (equivalent to t-values above 4.7; t-threshold when correcting for multiple comparisons is about 3.7). With this representation most of the sulci, with greenish color, show no difference between the two groups. No region shows large differences (there is no dark blue or red regions). Finally, the occipital lobe, the superior temporal sulcus and the polar frontal sulcus show some differences both in the right and left hemisphere.

**Figure 10.**
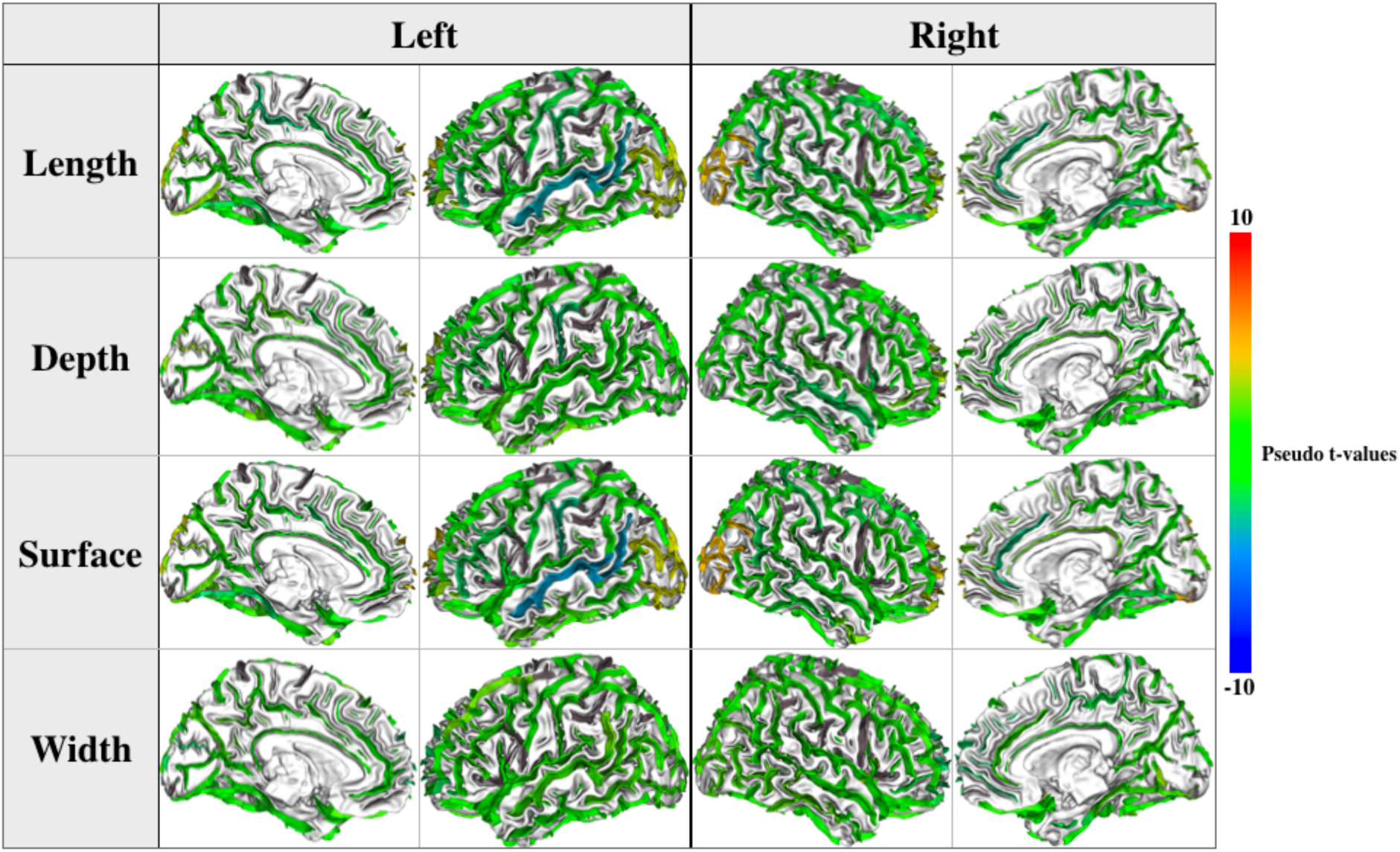
Pseudo T-values (square root of the F-score obtained in a linear model with age and sex as covariates) for the differences between the two groups for our four measures. Sulci with no values appear in grey.

In Figure 11 we report the individual values for the regions identified in Figure 10 as well as the 99.9% Confidence Intervals of the group difference (arbitrary correction for multiple comparisons; see methods). The results did not suggest any real difference between both groups.

**Figure 11.**
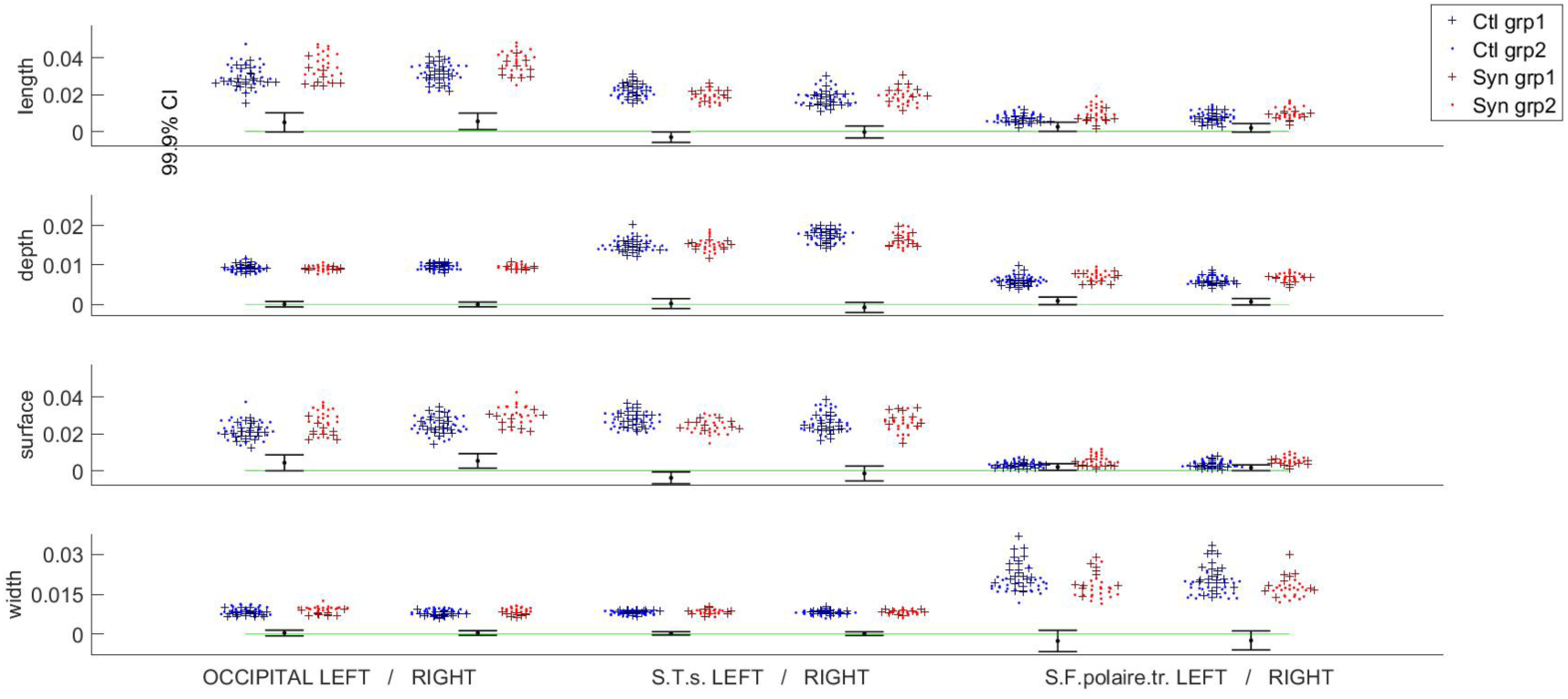
Sulcal length, mean depth, surface and width normalized for the total value in controls (blue) and synesthetes (red), in Study 1 (dark crosses) and 2 (bright dots) for three regions that show some differences. CIs were computed with age and sex as covariates (no interaction term). Only values for the most different sulci between the two groups were shown (occipital and parietal lobes).

## 4. Discussion

Several publications have reported structural brain alterations in synesthetes compared to controls. A recent review (3) shows a lack of consistency of the localization of the reported alterations and more importantly points out several methodological flaws. Our goal here was also to question the results we obtained in a recent study on graphemes-color synesthesia (8) and extend our study in introducing new features for anatomical comparison.

### 4.1 Replication of a VBM study

Some image processing steps may influence the quality of voxel based morphometry results, especially realignment and segmentation procedures (42-44). Firstly, we reanalyzed our 3T Bruker data with an updated implementation of the VBM method (VBM8) and the same general linear model. The results show (see Table 1 and Figure 1) that the differences initially observed in the RSC and STS were detected but were weak (notably, they did not reach the classical “statistical threshold” anymore). Clearly, the improvement of the segmentation algorithm has an impact on the robustness of the detected differences. A small difference in the fusiform gyrus was revealed (X=-26, Y=-72, Z=-12). Several authors reported this region as color-sensitive. For instance, the color-sensitive area was defined in human at -27 -57 -11 (40); -33, -65, -14 (38); -29 -68 -14 (39) and -26 -81 -9 (8) (see (8) for a detailed discussion about the “color center” localization). Change in this region has been considered as the main cause for color synesthesia (45, 46). However, recent attempts to find differences in this region were not convincing (see (3)). Our results (see Figure 2) reveal all structural differences including those not passing an arbitrary threshold and suggest that the screening of a larger synesthetes population could reveal significant differences.

For replication, we considered a new population of grapheme-color synesthetes (n= 21) and controls (n=26), and analyzed the 3T Philips data with VBM8 using the model previously used. The comparison of Figure 2 and Figure 4 indicates that potential structural differences were detected at different spatial locations based on either Study 1 or Study 2. When we pooled the two populations (32 vs 32) some differences detected in Study 1 (see Figure 5, Table 2) in the fusiform gyrus were still visible. Two different MR scanners were used and because the ratio of cases to control was different between the studies, a covariate was introduced in our model for these two different acquisition conditions (47). Pooling data from different scanners introduced confound in VBM analysis (48). Because the value of the magnetic field (3T) and the spatial resolution of the structural images were respectively identical, the “scanner effect” or “sequence” effect may be limited (49). Note that if the initially detected differences were “real” (i.e. not due to sampling noise), they would be likely detected with a new MR scanner generation equipped with a 32 channels head coil (Study 2). The extent of the confidence intervals for the group differences displayed in Figure 3 provides an idea of the order of magnitude of “true” differences compatible with our sample in Study 1, Study 2 and their pooling. There could be in fact no difference at all, but differences of up to 0.05 of WM probability are also compatible with our data. Such a difference (for example a change from 0.3 to 0.35) may be physiologically relevant, but could be assessed with confidence only by testing many more subjects.

### 4.2 Structural connectivity

In order to search for other possible structural differences in the two populations, we acquired Diffusion Tensor Imaging data. Using mean diffusivity (see Figure 6) we failed to detect differences. A local increase of Fractional Anisotropy would reveal more white matter and potentially more local connections (hyperconnectivity). Table 3 and Table 4 indicate respectively local increase and decrease in FA in the brain of synesthetes. Figure 7 shows increase in FA for some clusters. The confidence interval values for the group difference indicate that the measured differences were weak (Figure 8). Our results did not support previous claims using the same TBSS analysis procedure with (13, 14, 50) or without (15, 51) *a priori* hypothesis. Since the first attempt to detect structural differences in the brain of synesthetes using DTI (14), several improvements have been performed, from image acquisition (e.g. increase in the number of gradient directions), processing (e.g. geometric distortion correction) and analysis (e.g. crossing-fiber detection). A larger group study using state-of-the-art DTI acquisition and analysis procedures is needed for further structural connectivity investigation in synesthetes.

### 4.3 Sulcal shapes

Finally, using a recent sulci-based morphometry technique (34) we searched for differences at the sulcal level. We considered four features: length, depth, width and surface for 45 sulci (see Figure 9). Figure 10 shows all the measured differences for these features for 28 synesthetes versus 44 controls. It clearly appears that no important difference in cortical anatomies was found between controls and synesthetes. Only weak differences appear in the superior temporal sulcus and the occipital lobe. The former result may be in line with the positive correlation between fractional anisotropy in the right temporal cortex and scores of synesthesia strength reported in (14). The latter is in coherence with our VBM results of Study 1 (see Figure 2). Figure 11 indicates the corresponding confidence intervals. Our study reports the first investigation of sulcal shapes in synesthetes compared to controls, but due to large individual variations of sulcal anatomy, processing a large cohort is required to assess the possible relationship between brain folding and synesthetic experiences. For instance (33) involved 1009 healthy young adults for understanding genetic factors contributing to sulcal shape variability.

### 4.4 Shifting from NHST

To facilitate comparison with the literature we have reported p-value corrected for multiple comparison. Eklund et al. (17) demonstrated that multiple comparison correction procedures used in neuroimaging for cluster-wise inference artificially inflate false-positive rates. They considered that cluster failure for fMRI data inferences was mainly due to the false assumption of the gaussian shape of spatial autocorrelation functions. Similarly for VBM studies, the difficulty to control false positive is due to the non-normality of the data, even after spatial smoothing, and directly dependent on sample size (52, 53). More generally (16) underlined that multiple testing corrections do not prevent from false positive findings, especially in life science where studies are in general underpowered face to the large set of potentially influent variables difficult to master. In the same line, several authors have emphasized that null hypothesis significance testing (NHST), leading to the unsolvable problem of false positive control, is a theoretical framework maladapted to determine the significance of the results in social, psychological and cognitive sciences and should be banished (see the virulent attack from the statistician J. Cohen (54) or (55); see also (56) for MRI studies). A statistic paradigm shift is then proposed with ‘New Statistics’ (18) reporting effect sizes and confidence intervals rather than p-values or, for neuroimaging studies, statistical maps at a predefine p-value threshold. Several arguments support moving beyond NHST toward a cumulative quantitative approach where data are presented in a way that facilitates their interpretation in context. We promote such an approach in our paper with the dual-coding data visualization proposed by (19).

### 4.5 Data sharing

As reported by several authors (57, 58), low statistical power is endemic in Neurosciences. Our study clearly demonstrates that the use of a small-size population leads to unreproducible results even when inclusion criteria for group definition are strict and false-positive rate “controlled”. Data sharing between laboratories is the only way to improve statistical power in our neuroscience studies, increase reliability and confidence regarding effect size. This is more pregnant in domains, such as synaesthesia, where the recruitment of subjects is costly, long and time-consuming and the effect to measure, if any, is subtle, as evidenced in the present study. Knowledge about this subject will progress only if we decide to share our data. It is then imperative that all the steps of data processing be accurately reported and precise rules for statistical analysis respected (see (59, 60) for interesting recommendations). The ideal number of subjects for detecting a reproducible difference is not straightforward. Because the effect size may be low, more than one hundred individuals in each group seems a minimum (61). For going in this direction, all our structural data (T1-weighted and DTI) are freely available on request (https://shanoir.irisa.fr/Shanoir/login.seam, contact M. Dojat). Please refer to the present paper in case of the reuse of these datasets.

## Acknowledgments

We thank the technical staff at the 3T scanner facility from Grenoble MRI facility IRMaGe. IRMaGe was partly funded by the French program Investissement d'avenir run by the Agence Nationale pour la Recherche; grant Infrastructure d'avenir en Biologie Sante - ANR-11-INBS-0006. This project was funded in part by the Agence Nationale de la Recherche ANR-11-BSH2-010.

